# RAD-sequencing for estimating GRM-based heritability in the wild: a case study in roe deer

**DOI:** 10.1101/496083

**Authors:** L Gervais, C Perrier, M Bernard, J Merlet, J Pemberton, B Pujol, E Quéméré

## Abstract

Estimating the evolutionary potential of quantitative traits and reliably predicting responses to selection in wild populations are important challenges in evolutionary biology. The genomic revolution has opened up opportunities for measuring relatedness among individuals with precision, enabling pedigree-free estimation of trait heritabilities in wild populations. However, until now, most quantitative genetic studies based on a genomic relatedness matrix (GRM) have focused on long-term monitored populations for which traditional pedigrees were also available, and have often had access to knowledge of genome sequence and variability. Here, we investigated the potential of RAD-sequencing for estimating heritability in a free-ranging roe deer population for which no prior genomic resources were available. We propose a step-by-step analytical framework to optimize the quality and quantity of the genomic data and explore the impact of the SNP calling and filtering processes on the GRM structure and GRM-based heritability estimates. As expected, our results show that sequence coverage strongly affects the number of recovered loci, the genotyping error rate and the amount of missing data. Ultimately, this had little effect on heritability estimates and their standard errors, provided that the GRM was built from a minimum number of loci (above 7000). GRM-based heritability estimates thus appear robust to a moderate level of genotyping errors in the SNP dataset. We also showed that quality filters, such as the removal of low-frequency variants, affect the relatedness structure of the GRM, generating lower *h*^2^ estimates. Our work illustrates the huge potential of RAD-sequencing for estimating GRM-based heritability in virtually any natural population.

## Introduction

Estimating the evolutionary potential of quantitative traits and reliably predicting responses to selection in wild populations are important challenges in evolutionary biology. However, measuring the additive genetic variance and heritability of a trait, and the genetic correlation between a trait and fitness components (i.e. predicting the response to selection), require estimates of pairwise relatedness between individuals. For wild populations, these estimates have traditionally been based on a multigenerational pedigree. However, many species of research interest are hard to sample with the intensity and long-term effort required for pedigree construction (Pemberton, 2008). This has led to a strong taxonomic bias, with the great majority of heritability estimates available for vertebrates, especially birds and large mammals, because of their ease of monitoring (Postma, 2014).

Advances in high-throughput sequencing technology have opened up the possibility of getting access to the realized proportion of the genome that is shared among individuals (i.e. generating a Genomic Relatedness Matrix - GRM) in virtually any non-model species, with the potential to greatly expand the taxonomic coverage of quantitative genetic studies in the wild (Gienapp et al., 2017). While first applied to human data (Yang et al., 2011), GRM have recently been shown to provide heritability estimates which are similar to those from multigenerational pedigrees in free-ranging animal populations (Bérénos et al., 2014 on Soay sheep, Robinson et al., 2013 on great tits, Perrier et al., 2018 on blue tits). These studies have demonstrated that GRMs computed from a few hundreds of individuals and a few thousand loci may be enough to obtain heritability estimates with low standard interval errors when effective population size is relatively small and linkage disequilibrium (LD) high (Bérénos et al., 2014; Perrier et al., 2018; Stanton-Geddes, Yoder, Briskine, Young, & Tiffin, 2013).

The few studies that have attempted to estimate GRM-based heritabilities in the wild have all focused on long-term monitored populations for which pedigrees were already available. Moreover, the authors often had access to knowledge about both genome sequence and variability on the study species, or on a closely related species, facilitating the development of the genotyping SNP array. Whole genome sequencing is still prohibitive for many species in terms of cost, bioinformatic resources especially for organisms with large genomes, and required DNA quality (Ekblom & Wolf, 2014). Recently, a variety of Restriction site-Associated DNA sequencing (RAD-seq - Andrews et al 2016) methods and other similar approaches to sequencing a subset of the genome have been developed. These approaches are increasingly attractive because they are cost-effective, not dependent on prior genomic information and SNP loci are discovered and genotyped in a single procedure (Narum, Buerkle, Davey, Miller, & Hohenlohe, 2013), making them applicable to a wide range of non-model species. RAD-seq approaches are also valued for their high level of flexibility, allowing optimization of the trade-off between the number of markers, the number of genotyped individuals and sequencing depth in an Illumina run using a wide variety of experimental designs and using different restriction enzymes (Peterson, Weber, Kay, Fisher, & Hoekstra, 2012).

Despite the numerous benefits, the potential of RAD-seq for estimating GRM and quantitative genetic parameters in the wild, particularly in non-model species, has received little attention (but see Perrier et al., 2018). One reason is that long-term research projects willing to genotype thousands of individuals to estimate GRM often invest in developing a SNP chip. Another reason is that RAD-seq approaches are reputed to lead to high rates of missing data and allele dropout that might bias relatedness estimates and downstream biological inferences (Dodds et al., 2015; Gienapp et al., 2017). Furthermore, although easy-to-use bioinformatic pipelines are available for the *de novo* assembly of loci and SNP genotyping (J. M. Catchen et al., 2017; Eaton, 2014), little is known about the impact of sequencing strategy (e.g. marker read-depth coverage) and parameter choice during the SNP calling/filtering process on genotyping error/missing data rates, GRM structure and, ultimately, quantitative genetic estimates. Several recent studies have proposed methodologies to optimize the *de novo* assembly of markers and minimize error rates (Fountain, Pauli, Reid, Palsbøll, & Peery, 2016; Mastretta-Yanes et al., 2015; Paris, Stevens, & Catchen, 2017; Rochette & Catchen, 2017). Others have explored how relatedness estimators can circumvent the influence of genotyping errors (Attard, Beheregaray, & Möller, 2018) or tested the influence of the number of markers/samples on heritability estimates from whole genome data (Stanton-Geddes et al., 2013). However, to our knowledge, no study has explicitly explored the potential limits and pitfalls of RAD-seq for estimating quantitative genetic parameters in the wild.

In this study, we describe an analytical framework (see Figure 1) to estimate trait heritability from RAD sequencing data, using a free-ranging roe deer population (*Capreolus capreolus*, Linnaeus 1758) as a case study. We focus on the heritability of body mass, a trait closely associated with both survival and reproductive performance (Hamel, Gaillard, Festa-Bianchet, & Côté, 2009; Quéméré et al., 2018). There are three key parameters for estimating GRM-based heritability: the number of SNPs, the number of individuals and sampling variance in relatedness (Visscher & Goddard, 2015). First, we provide guidance on the sampling and sequencing strategy. Using *in silico* simulation and double-digest RAD-sequencing (Peterson et al., 2012), we established a sequencing strategy that optimizes the balance between the number of loci and the number of individuals genotyped, and library/sequencing costs. The sequencing depth (i.e. average read depth per locus) may directly affect data quality, with potential impact on biological inferences (Sims et al 2014). We, therefore, explored how variable sequencing effort (targeted coverage depth of 20x *versus* 60x) affects the rates of genotyping error and missing data, the accuracy of relatedness coefficients and, ultimately, heritability estimates. We then detail the different steps in the bioinformatic and analytical pipeline from the raw sequence data to the implementation of the GRM and the estimation of GRM-based heritability. Specifically, we explore how parameter choice during the *de novo* assembly of loci and the SNP data filtering may influence data quality (genotyping error rate) and quantity (number of informative loci), the structure of the GRM and GRM-based heritability. Lastly, we provide practical recommendations to minimize bias in the estimation of relatedness while maximizing the explanatory power of the GRM.

**Figure 1.**
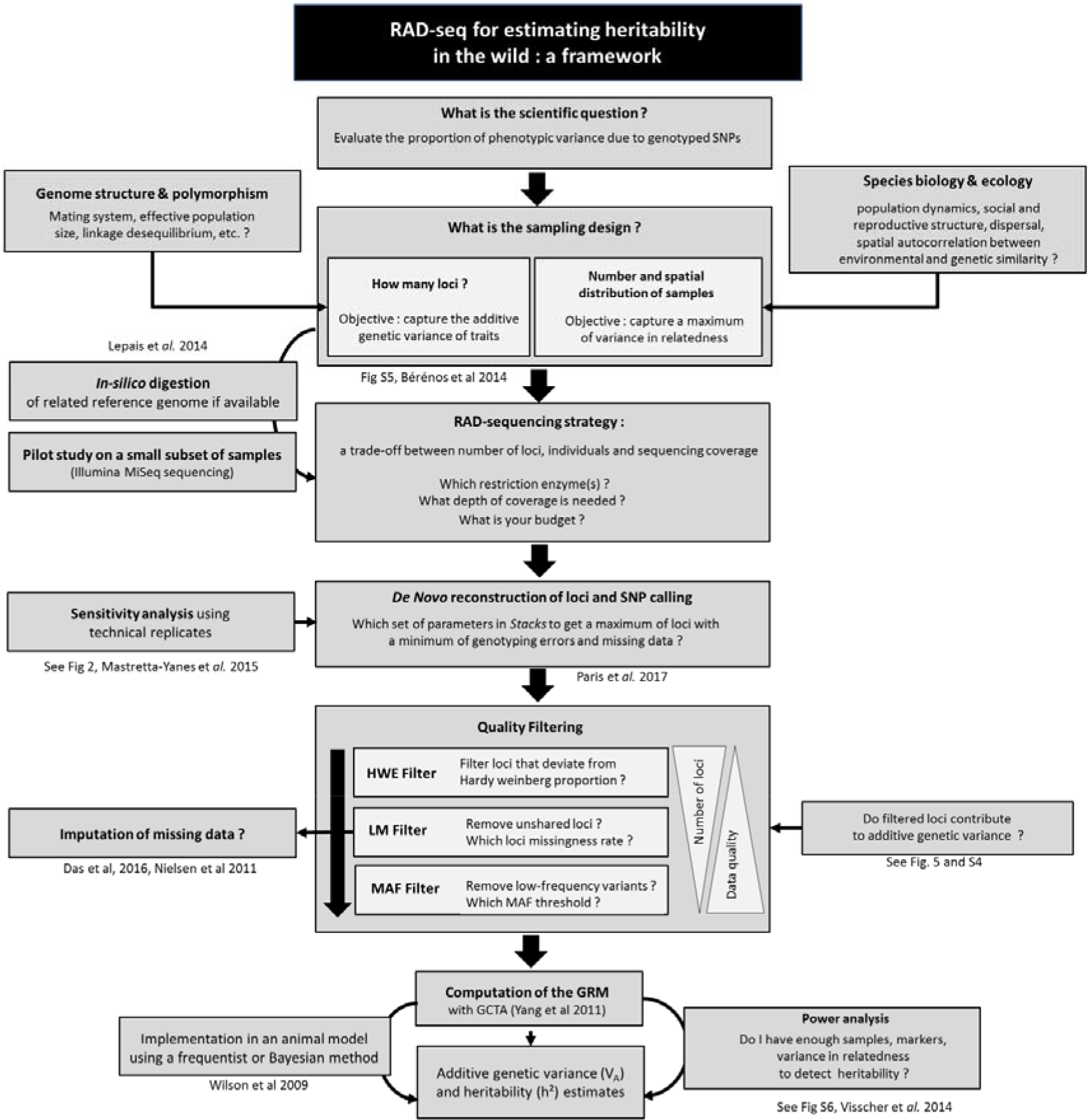
Practical framework with steps for estimating a GRM-based heritability from RAD-seq data. Throughout the process (sampling design, SNP calling, quality filtering), there are trade-offs between data quality (choice of sequencing coverage, genotyping error/ missing data rates) and quantity (number of samples and loci).

## 2. Methods

### 2.1. Sampling design

The study focused on a roe deer population inhabiting a heterogeneous agricultural landscape in southwestern France (43°13HN, 0°52HE). We selected a set of 249 individuals which were caught using large scale drives during winter, between 2002 and 2016 at seven sampling sites, separated by a few kilometers, and with a variable proportion of woodland vs. open areas (e.g. crops, meadows, pasture) (Figure S1). Adult roe deer generally occupy a single home range across their entire adult life, whereas around 40% of juveniles disperse (Debeffe et al., 2012). Our hierarchical sampling scheme is thus expected to maximize variance in relatedness by including potentially closely related individuals within sampling sites, individuals with intermediate relatedness among sampling sites and unrelated individuals that immigrated from outside the study area. At each capture, animals were sexed and weighed. Age was either known (for individuals first caught as newborns or juveniles) based on tooth eruption, or estimated from tooth wear (for individuals first caught older than one year). All applicable institutional and European guidelines for the care and use of animals were followed. All the procedures involving animals were approved by the Ethical Committee 115 of Toulouse and authorized by the French Ministry in charge of ethical evaluation (n° APAFIS#7880-2016120209523619v5).

### 2.2. Sequencing strategy

DNA was extracted from skin samples using DNeasy™ Tissue Kit (Qiagen). SNP genotyping was performed using double-digest restriction site associated DNA sequencing (ddRAD-seq, Peterson et al., 2012). This genotyping-by-sequencing method was chosen because it offers greater flexibility than traditional RAD-seq, with the possibility of optimising the number of loci sequenced by testing different combinations of restriction enzymes and fragment size selection. This optimization process was carried out by performing *in-silico* digestions of the red deer genome (*Cervus elaphus*) (Bana et al., 2018) using the R package Sim-RAD (Lepais & Weir, 2014). We tested various experimental designs and finally retained the two enzymes *EcoR1* and *MsPI* and size-selected libraries of 270-330 bp insert-size that produced around 35,000 fragments in the red deer genome. Using this design, we expected to sequence 96 individuals with an average of 60X coverage in a single Illumina HiSeq 2500 lane (based on average read counts of 200 M reads per lane). To ensure that this design provides at least 15,000 polymorphic loci (i.e. the minimum required to capture the additive genetic variance of traits in Soay sheep Bérénos et al., 2014), we performed a pilot study on a set of 12 samples sequenced using an Illumina MiSeq lane. We then applied our protocol to the whole dataset of 249 samples which were multiplexed in equimolar proportions in groups of 24 individuals and sequenced on Illumina HiSeq 2500 (V4) (Paired-end reads of 125 bp). Library- preparation and Illumina sequencing was performed at the NERC Biomolecular Analysis Facility-Edinburgh (Genepool, Edinburgh, UK). To evaluate the reliability of the genotyping process and optimize the *de novo* loci assembly (see below), seven individuals were repeat-processed, either at the library preparation step (3 pairs of ‘library replicates’ individuals) or at the sequencing step (4 pairs of ‘sequencing replicates’).

### 2.3. De novo reconstruction of loci and error quantification

Raw sequences were first inspected with FASTQC (Andrews 2010) for quality control. Reads were then demultiplexed (i.e. assigned to each sample) and trimmed to 117 bp using *process_radtags* (part of the Stacks 1.35 pipeline, Catchen et al. 2013) without allowing any mismatch in the barcode sequence. The *‘de novo map’* pipeline of Stacks was used to build loci *de novo* (without a reference genome) and call single nucleotide polymorphisms (SNPs). *De novo* loci assembly in Stacks is governed by three main parameters that influence the balance between the number of loci, genotyping errors and missing data. The first two parameters affect the way loci are built at the individual level: *-m* is the minimum number of reads to form a stack (allele) and *-M* is the maximum number of mismatches allowed between alleles. A high value of –*m* may cause allele dropout, while a low value may generate false alleles (Catchen et al., 2013). A high value of –*M* may generate false heterozygotes (due to the erroneous combination of different homozygous loci), while a low value may erroneously generate homozygous loci. The third parameter -*n* is the maximum number of mismatches allowed between homologous loci across all samples to build the population catalog.

The choice of parameter values is specific to each study since it depends on the biology of the study species (inherent polymorphism, ploidy level) and the experimental design (restriction enzyme used, number of samples multiplexed, sequencing coverage) (Paris et al., 2017). Following Mastretta-Yanes et al., 2015, we carried out a preliminary analysis to identify the parameter values that maximized the number of loci recovered while minimizing genotyping error rates in our case study. To guide our choice, we estimated two error rates between the seven pairs of technical replicates: (1) the *locus error rate* (LE_R_) corresponding to the number of loci present in only one of the two replicates, divided by the total number of loci being compared (i.e. missing data at the locus level) and (2) the *allele error rate* (AE_R_), calculated as the number of incongruent genotypes between the two replicates, divided by the number of common loci. We first explored the influence of each parameter (-m, -M and –n) one-by-one (while holding the two others fixed) (see Text S1 in supplementary material). On this basis, we tested four sets of parameters corresponding to different ways to deal with the trade-off between data quantity and quality: (S1) parameters designed to maximize the number of markers (‘MaxLoci’, m=2, M=2, n=1), (S2) parameters designed to minimize the error rates (‘MinError’, m=11, M=2, n=1), (S3) parameters designed to lead to a low error rate with an intermediate number of markers (‘Intermediate’, m=7, M=2, n=1) and (S4) parameters by default (‘Default’ hereafter, m=3, M=2, n=0). This preliminary analysis was restricted to loci with a single SNP that was shared by at least 80% of individuals. We retained the parameter combination that provided the optimal balance between number of loci and genotyping error and used it on the full dataset (249 samples).

### 2.4. Quality filtering

Once the SNP calling process is completed, additional filtering steps are regularly performed to remove loci and/or individuals with too many missing data (loci and individual missingness rates respectively) and persistent genotyping errors (removing rare variants and loci that deviate from Hardy-Weinberg equilibrium). Here, we explored the impact of these quality filters on genotyping and locus error rates, number of loci and GRM structure (relationship coefficients and overall variance in relatedness). We successively applied three filters to the SNP dataset by testing different threshold values using VCFtools (Danecek et al. 2011): (1) the “HWE filter” removes loci that deviate from Hardy-Weinberg Equilibrium (under the assumption that the population is panmictic) (P<0.05) (2) the “LM filter” (Loci Missingness) discards loci with a missingness rate above 10%, 20%, 30% or 40% and (3) the “MAF” filters loci with Minimum Allele Frequency greater than 1%, 5% or 10%. We retained one SNP per locus (to minimize linkage disequilibrium between markers) and removed individuals with more than 50% missing data. All analyses were performed on the dataset built with the S3 “Intermediate” *stacks* model described above which offered the best compromise between the number of loci and genotyping error rate (see results below).

### 2.5. Inference of GRM-based heritability

GRMs were computed using GCTA from identify by state (IBS) SNP relationships (Yang et al 2011). At each locus, relatedness was scaled by the expected heterozygosity 2pq (Yang et al. 2010, 2011). GRMs were then fitted in a REML-mixed-linear model to estimate the amount of phenotypic variance in body mass that was explained by additive genetic variation (SNPs). Univariate animal models were run in ASREML-R v3.00 (VSN International, Hemel Hempstead, UK) as follows:

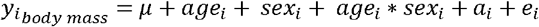

where *μ* is the *population* mean, *a_i_* is the breeding value (i.e. effects of i’s genoptype relative to *μ*) and *e_i_* the residual term. The individual’ s sex and age as well as the interaction between them (*age_i_*sex_i_*) were included in the model as fixed effects (Gaillard, Delorme, & Jullien, 1993). Narrow-sense heritability of body mass was estimated as h^2^ = V_A_/V_P_ where V_A_ is the additive genetic variance and V_P_ the total phenotypic variance (V_A +_ V_R_). Residuals and fitted values were visually inspected to verify model assumptions. We investigated the influence of SNP calling (parameters in *Stacks*) and quality filtering (LM and MAF filter) on GRM-based heritability estimates. First, we explored the influence of loci missingness rate (LM ranging from 10% to 40%) by fixing the MAF to 1%. Then, we evaluated the impact of MAF (1%, 5% or 10%) by fixing the LM ratio to 20%. These analyses were carried out on SNP datasets called with the four sets of parameters in Stacks (‘Max Loci’, ‘Min Loci’, ‘Intermediate’ and ‘Default’). Since these four models generated a variable number of informative loci, and because estimates of heritability increase up to an asymptote with the number of SN Ps used to build the GRM (Bérénos et al., 2014; Perrier et al., 2018), to be comparable, the *h^2^* value must be estimated using GRMs built from the same number of SNPs. Here, we fixed the number of SNPs to the minimum obtained in all analyses and performed fifty resampling iterations.

### 2.6. Influence of sequencing coverage

Genotyping error rates are assumed to be highly sensitive to sequencing coverage, particularly in *de novo*-assembled data sets (Catchen et al., 2013; Fountain et al., 2016). To explore the effect of sequencing coverage, we randomly sampled 20% of the raw reads of the initial dataset (using Seqtk: https://github.com/lh3/seqtk) so as to obtain an expected average read depth of 15 reads per locus (20x, hereafter ‘low-coverage dataset’). We then calculated genotyping error rates (LE_R_ et AE_R_) between pair of replicates as we did for the full dataset and used the same sensitivity analysis described above to select the parameters in *Stacks* that offered the best compromise between data quality and quantity. Then, we evaluated how the filters affected the number of polymorphic loci, relationship coefficients, variance in relatedness of the GRM and heritability estimates.

## 3. Results

### 3.1. Optimization of the de novo loci assembly of loci and SNP discovery

An average of 6.9 million reads per individual was obtained after demultiplexing. The number of polymorphic loci recovered by *Stacks* in the exploratory analysis ranged from 14,536 for the ‘MinError’ parameter set to 21,681 loci for ‘MaxLoci’, with a median depth coverage per individual and per locus of between 39.1 and 88.4 reads. Median locus error rate (LE_R_) across the seven pairs of replicates varied between 3.1% for ‘MaxLoci’ and 5.1% for ‘default’ (Figure 2). Median allele error rates (AE_R_) ranged from 1.1% for ‘MinError’ to 3.2% for ‘MaxLoci’. We retained the parameters that offered the optimal balance between data quantity and quality for further analysis: ‘MinError’ and ‘Intermediate’ models generated very similarly low error rates, but the ‘intermediate’ model yielded more loci (17,013) and less variation in LE_R_ across the seven pairs of replicates. When loci assembly was attempted using a subset of our data (20% of the reads), loci and allele error rates were two to three times higher than when using the original dataset (Median LE_R_ ranging from 4.8% to 16%; Median AE_R_from 1.8% and 7.6%, Figure S2). The number of loci recovered also dropped sharply from 17,534 polymorphic loci for ‘MaxLoci’ to 2,092 for the ‘MinError’ model.

**Figure 2.**
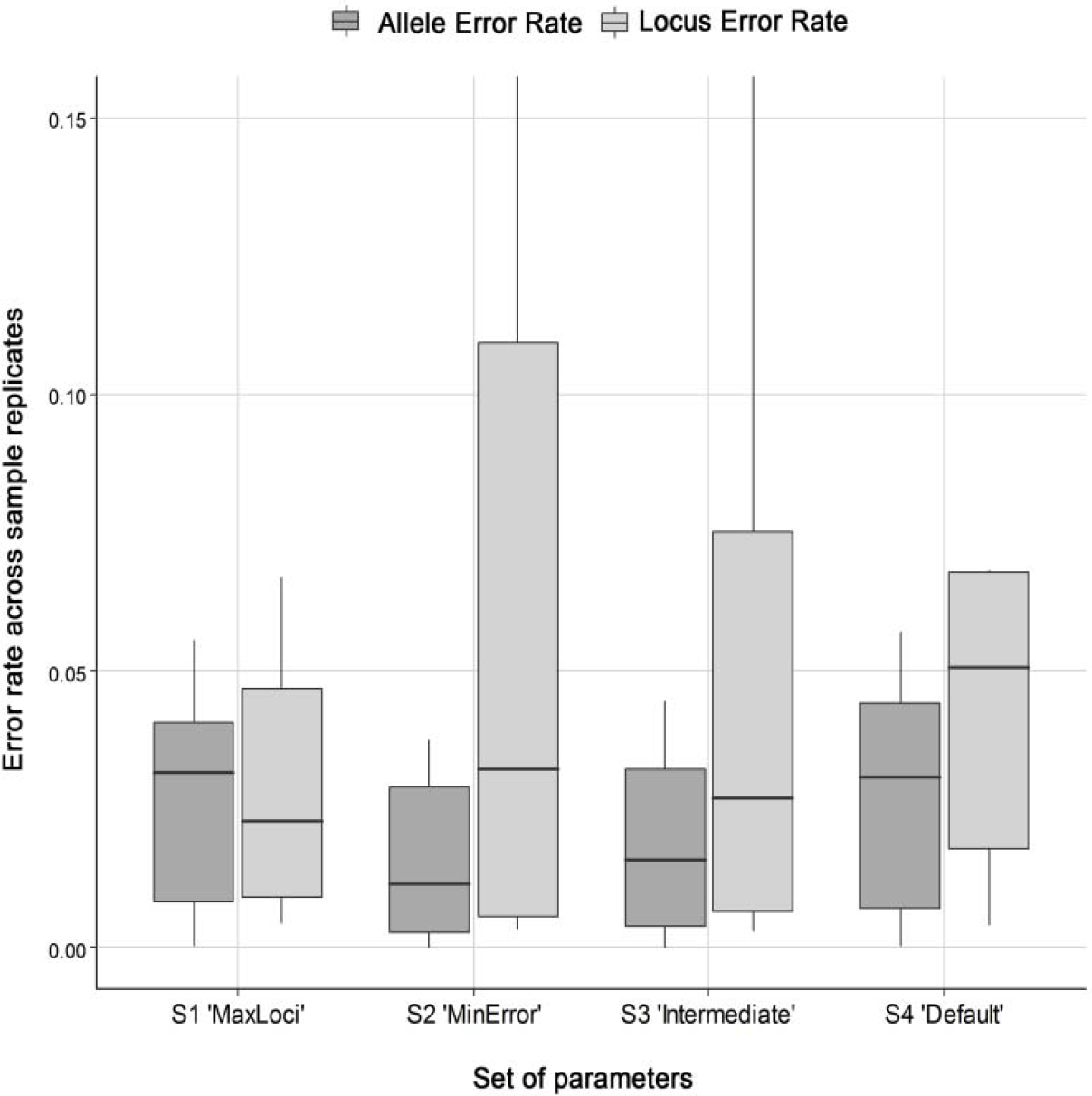
Effect of the different sets of parameters in *Stacks* on allele and locus error rates for the high-coverage dataset. (error rate distribution across the 7 replicate pairs). For each *Stacks* profile and each pair or technical replicates, we computed the locus error rate (LE_R_) corresponding to the number of loci present in only one of the two replicates, divided by the total number of loci being compared (i.e. missing data at the locus level) and the allele error rate (AE_R_), calculated as the number of incongruent genotypes between the two replicates, divided by the number of common loci.

### 3.2. Quality filtering: impact on genotyping error rates, number of loci and GRM

#### HWE filter

Once the *de novo* assembly of loci had been completed on the full dataset (249 samples, using the ‘intermediate’ stacks model), we obtained 96,773 loci totaling 154,540 SNPs. When removing loci that deviated from HWE, we retained 83,893 SNPs (Fig 3a). The ‘HWE filter’ had very little impact on loci and allele error rates (Fig 3b,c), but led to a substantial decrease in the number of SNPs (-13%) and the off-diagonal variance in relatedness of the GRM (V_GRM_ = 0.6 x 10^-3^ *versus* 0.8 x 10^-3^ for the filtered and unfiltered datasets, respectively) (Fig 3d).

**Figure 3.**
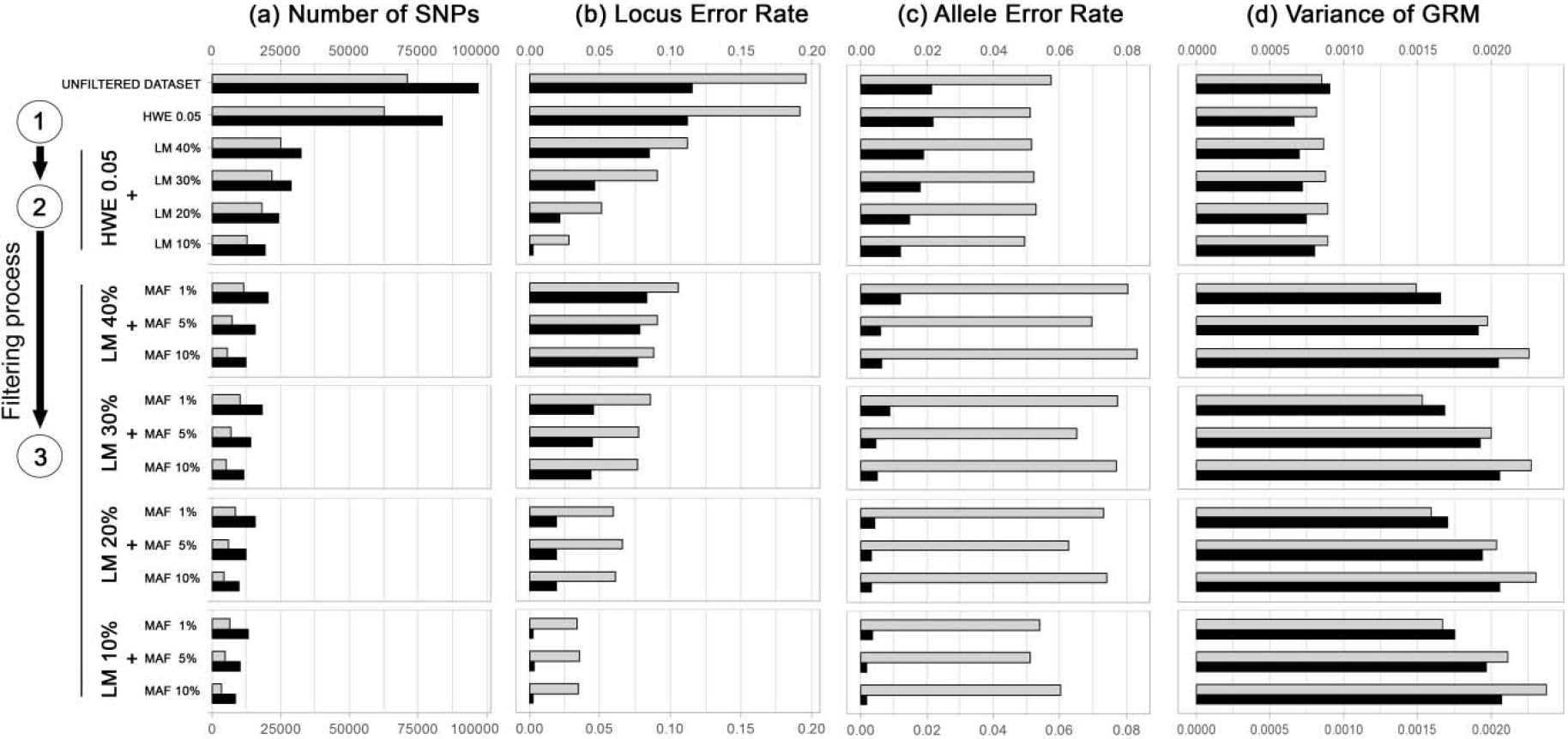
Impact of quality filtering on genotyping error rates, number of markers and GRM variance. Evolution of the (a) number of SNPs, (b) median locus error rate and (c) allele error rates (across replicates) and (d) the variance in relatedness of the GRM (V_GRM_), as a function of the filters successively applied for the low-coverage (in grey) and high-coverage (in black) dataset: We first removed loci that deviated from Hardy-Weinberg equilibrium (P<0.05) (step 1). We then discarded loci with missingness rate >10%, 20%, 30% or 40% (step 2). Lastly, we filtered loci with a minimum allele frequency (MAF) greater than 1%, 5% or 10% (step 3).

#### Loci missingness filter (LM)

When we applied the ‘loci missingness’ filter (LM), we observed a two to three-fold decrease in the number of loci: between 19,493 and 32,538 SNPs were retained when removing loci typed in less than 90% (LM=10%) and 60% (LM=40%) of individuals respectively. Variance in relatedness did not change markedly (V_GRM_ ranging from 0.6-0.7 x 10^-3^), but the remaining loci had significantly lower error rates, particularly with a LM = 10% (LE_R_ ~ 0.03%, AE_R_ ~ 0.02%).

#### MAF filter

Filtering rare variants only affected locus error rate slightly, but this resulted in a lower allele error rate, particularly when using high LM: AE_R_ fell from 1.9% to 0.6% when using LM 40% and MAF 10% filters. However, this stringent MAF threshold also led to a >50% decrease in the number of markers (to a final number ranging between 8434 and 12528 SNPs). MAF filtering had also a marked influence on the GRM off-diagonal variance in relatedness (V_GRM_) which substantially increased from ~0.8 x 10^-3^ to ~1.6 x 10^-3^ when using a MAF=1% filter and to ~2 x 10^-3^ with MAF=10%. However, the MAF threshold had little effect on pairwise relatedness coefficients *per se*, although the highest coefficients tended to increase when using a stringent MAF filter (MAF=10%, Fig 4a).

**Figure 4.**
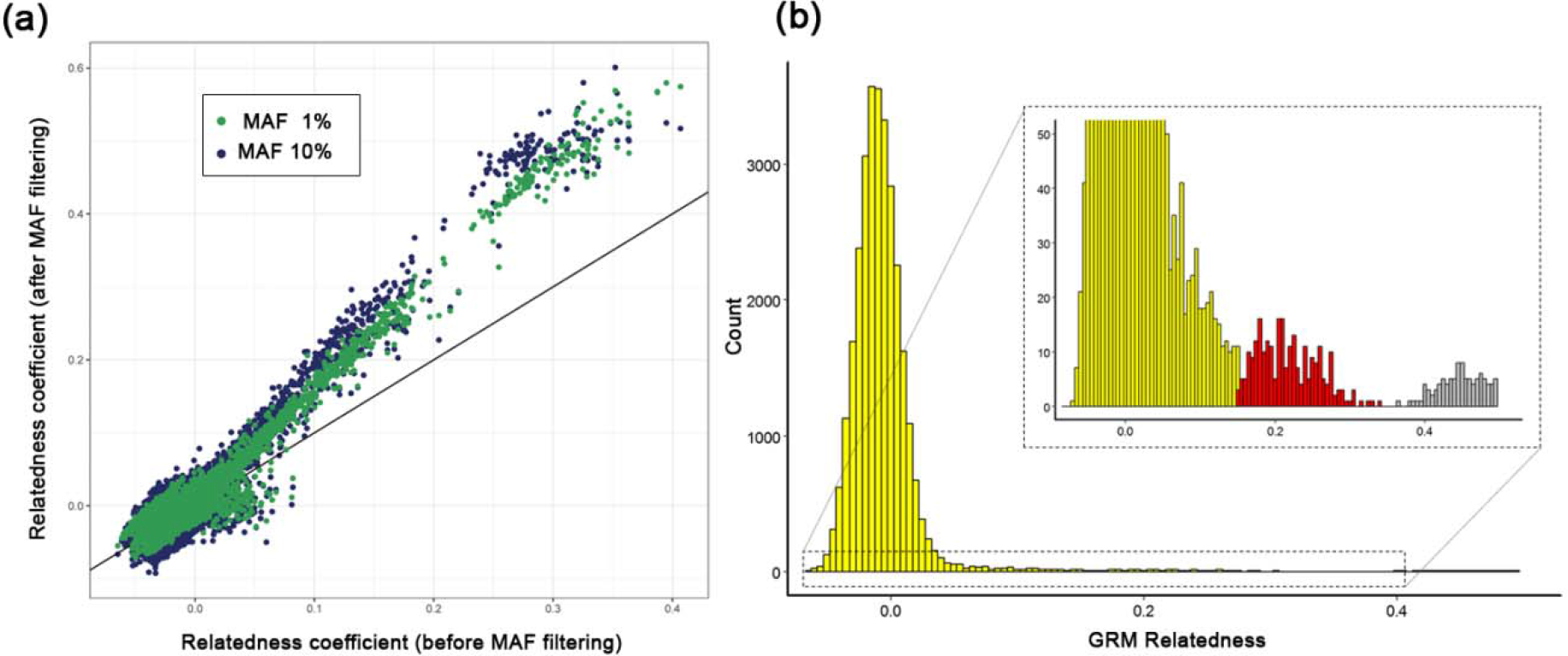
Genomic relatedness. (a) Biplot of genome-wide relatedness matrices (GRM) computed before and after filtering SNPS with a MAF 1% (green dots) or MAF 10% (blue dots) threshold (from the high-coverage dataset): The highest relatedness coefficients increased when applying a stringent MAF (10%). (b) Histogram of pairwise genome wide relatedness (using MAF=1% and LM=20%).

In comparison, the ‘low-coverage dataset’ provided 70,956 loci totaling 111,803 SNPs (i.e. a 27% decrease compared to the full coverage dataset) using the S4 ‘Default’ stack model (optimal model for this dataset). The number of loci fell to between 3,493 loci (LM=10%, MAF=10%) and 11,441 (LM=40%, MAF=1%) after the filtering procedure (Fig 3b). The LM filter substantially reduced locus error rate (from 19.6% to ~3.5%). However, even after using all filters, allele error rate remained 10 times higher than in the high coverage dataset. As in the full dataset, variance in relatedness markedly increased when using the MAF filter to reach values between 1.4 x 10^-3^ (LM=40%, MAF=1%) and 2.3 x 10^-3^ (LM=10%, MAF=10%). Modifying the MAF threshold led to more changes in relationships than with the high-coverage dataset (Pearson r=0.969, P<10^-16^) (Fig S3a): relatedness coefficients >0.1 tended to be higher with MAF=10% than with MAF=1%. Lastly, the full dataset tended to generate higher relatedness coefficients than that the low-coverage dataset (Wilcoxon signed-rank test, P<10^-7^, Fig S3b), probably because of a much lower genotyping error.

#### 3.3. GRM structure and GRM-based heritability

The GRM mostly included unrelated individuals (Fig. 4b in yellow). However, on closer inspection, the histogram of relatedness indicated the presence of related individuals made up of parent-offspring or full-sib links (grey) and a number of half-sib-like links with relatedness around ~0.25 (red). Estimated heritability of body mass (*h^2^*) ranged from 0.56 (se=0.14) (‘Default’ *stacks* model, LM 10%, MAF 10%) to 0.68 (se=0.15) (‘Default’ model, LM 30%, MAF 1%). The lowest *h^2^* values were obtained using MAF 10% and LM = 10% (Figure 5a, S4a) which correponds to the filtering parameters that yielded the smallest number of SNPs. As expected, *h^2^* gradually increased with the number of markers up to an asymptote (here around 7000-8000 SNPs) (Figure S5). However, even when the number of loci used to build the GRM was fixed, *h^2^* tended to be lower with a stringent MAF threshod (10%) (Figure 4b).

**Figure 5.**
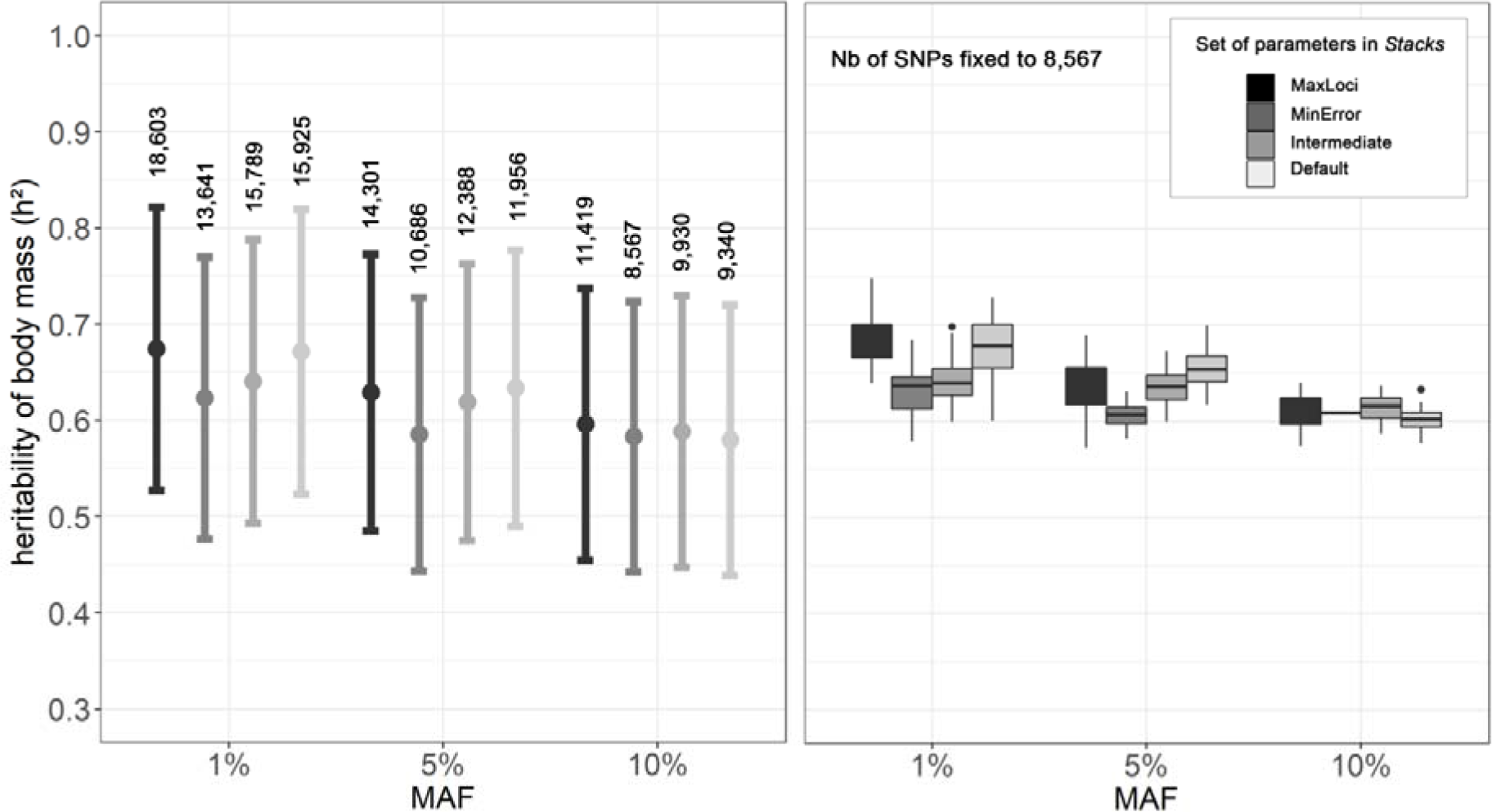
Sensitivity of estimates for h^2^ of body mass to SNP calling (*Stacks* model) and MAF threshold (high-coverage dataset): (a) raw *h^2^* estimates with standard error (error bars) obtained using GRMs computed with all SNPs generated by a given SNP calling (Stacks model) and filtering method (MAF = 1%, 5% or 10% and LM=20%). For each analysis, we reported the number of SNPs used to build the GRMs (b) *h^2^* estimates obtained by fixing the number of SNPs to 8,567 SNPs for all GRMs and resampling the SNP data 50 times. Boxplot and whiskers indicate median *h^2^* and variation across the 50 resampled datasets.

When we built the GRM from the low-coverage dataset using the same filtering procedure as with the full dataset (HWE, LM 10% and MAF 1%), we obtained *h^2^* estimates that ranged from 0.39 (se=0.13) (S2 ‘MinError’ stacks models, 216 individuals, 599 loci) to 0.67 (se=0.16) (S1 ‘MaxLoci’ *stacks* model, 246 individuals, 10,876 loci). In this case, the ‘MinError’ *stacks* model with a very high – *m* value (11) was not suitable as it removed too many loci and individuals (individuals with >50% missing loci). However, the three other models provided *h^2^* estimates that were similar to those produced from the full dataset, despite a lower number of loci (5,302 for S3 ‘Intermediate’, 8,646 for S4 ‘default’) and higher genotyping error rate (see above).

## 4. Discussion

Our results illustrate how RAD-sequencing data can be used to obtain relatively robust estimates of heritability in a free-ranging population for which no genomic resources are available. We showed that this process is not straightforward and that critical decisions must be taken during the sampling design and the computational process to optimize the quality and quantity of the genomic data. We showed that the choice of sequencing depth and of the values for bioinformatic parameters (calling and filtering genomic markers) markedly affect the rates of missing data and genotyping error, ultimately impacting the accuracy of genome-wide relatedness estimates. However, in our case study, these decisions had little impact on *h^2^* estimates, provided that the GRMs were built from a minimum number of loci (above 7000-8000 loci). This suggests that GRM-based heritability estimates are relatively robust to genotyping error and missing data in the SNP dataset. One important exception is the removal of rare variants that led to lower *h^2^* estimates when a high (10%) MAF threshold was used. We hereafter discuss the implications of our results and provide some practical recommendations for each step of the analytical process.

### 4.1 Sequencing design: target a high average sequencing coverage

The sequencing strategy, and especially read depth, may have a significant impact on the quality and quantity of the genomic data, especially when loci are built *de novo* i.e. without a reference genome (Catchen et al., 2013). The *in-silico* digestion of a related reference genome (red deer) followed by a pilot study on a small subset of samples enabled us to optimize the genotyping of around 70,000 loci with a mean coverage of 60 reads per locus (60x). Targeting such high sequencing coverage has multiple benefits. First, it allows maximization of the number of genotyped individuals. Indeed, it is frequent to observe among-sample variation in DNA quality and quantity during the RAD library preparation that may translate into variation in sequencing depth. When using a low average sequencing depth, individuals that are not sufficiently covered may be filtered out because they do not share enough loci with other individuals. Second, using a high average sequencing depth allows optimization of the number of loci and considerably reduces the rate of missing data and genotyping error. In our case, we obtained very low error rates with the 60x coverage dataset (LE_R_and AE_R_ ~ 2-3%, before filtering), well below the values obtained when genotyping was performed on a subset of our sequencing data (coverage ~20x, LE_R_ > 5% and AE_R_~2-8%). These error rates were also lower than those obtained by Mastretta-Yanes et al., 2015 *in Betula alpina* with a coverage ~ 10x (where LE_R_ >10% and AE_R_ > 5% regardless of assembly parameters). At the end of the SNP calling and filtering optimization procedure (using a MAF=1% and a LM=20%), we obtained nearly twice the number of SNPs with the 60x-coverage than with the 20x-coverage dataset (15,930 *versus* 8,809) with far fewer missing data (4.7% *versus* 7%). Several genotype imputation methods have been developed for handling missing data (Das et al., 2016; Nielsen, Paul, Albrechtsen, & Song, 2011), but incorrect SNP calling and allelic dropout are hard to detect and may have a strong impact on downstream population genetics and biological inferences (e.g. parentage assignment in Fountain et al.,2016; demographic inference in Shafer et al., 2017). In the past, because of budget limitations, sequencing coverage was often traded-off against increasing the number of individuals and loci genotyped (Mastretta-Yanes et al., 2015). However, the continuous improvement of Next Generation Sequencing technology has led to a rapid decrease in sequencing costs so that it is now possible to genotype tens of thousands of loci with >50x coverage for less than 50$ per sample (including library preparation cost). Hence, we strongly encourage targeting a high sequencing depth to maximize the quality and completeness of genomic datasets.

### 4.2. Impact of SNP genotyping and filtering on GRM and GRM-based heritability

Calling SNPs from RAD-seq data without using a reference genome to guide the assembly of loci is not a simple task since it requires the researcher to select the values for the bioinformatic parameters that determine how closely the sequences must match and the minimum coverage to identify true loci and alleles (Catchen et al., 2013; Paris et al., 2017). Our work illustrates the importance of performing a sensitivity analysis based on technical replicates to help select the optimal parameters and understand the limitations of the dataset (Flanagan, Forester, Latch, Aitken, & Hoban, 2018; Mastretta-Yanes et al., 2015). In our case, the high sequencing depth allowed us to design an assembly model with a relatively high minimum coverage threshold (7 reads) to recover allelic variants. This enabled us to eliminate most sequencing errors (reduced to <2%) while retaining a high number of loci with a minimum of missing data. However, for this dataset, genotyping error rates remained low (<5%) irrespective of the assembly parameters so that the SNP calling procedure ultimately had little impact on the mean and standard error of the body mass heritability estimate.

Additional quality control filtering steps are generally performed to further clean the dataset of uninformative markers and statistical artefacts based on loci missingness ratio, minor allele frequency and/or loci deviating from Hardy-Weinberg proportions (Huang & Knowles, 2014). The relevance of such filters is increasingly questioned, particularly when the final purpose is to estimate genomic relationships (Eynard, Windig, Leroy, Van Binsbergen, & Calus, 2015) or detect loci potentially under selection (Benestan et al., 2016; Roesti, Salzburger, & Berner, 2012; Waples, 2014). We explored the impact of each of these filters on the number of loci, error rates, GRM structure and GRM-based heritability. We found that the “HWE filter” had little impact on error rates, but led to a substantial decrease in the number SNPs and in the variance in relatedness of the GRM, two parameters that are critical for the estimation of GRM-based heritability (Bérénos et al., 2014). This filtering step is applied to remove sequencing or SNP calling error when the population is assumed to be panmictic. However, genotyping errors generally lead to only slight departures from HW proportions (Cox and Kraft 2006) in contrast to factors such as selection, age structure or non-random sampling that are often neglected (see Waples et al 2014 for a review). One can question the relevance of a filter that has little effect on the quality of the dataset but removes numerous loci that are potentially important for downstream analyses. In contrast, the “Loci missingness” filter (LM) appeared to have a significant impact on locus error rate that was reduced five-fold when excluding loci shared by less than 80% of individuals. Filtering unshared loci thus appears important to remove erroneous loci that are based on artefactual sequences during *de novo loci* assembly.

Lastly, as expected, we found that removing rare variants decreased allele error rate, particularly when a high MAF threshold was used (MAF < 10%). However, such a stringent MAF threshold also significantly reduced the number of loci and led to bias in the allele frequency spectrum. One of the major advantages of RAD-seq genotyping is that it captures rare variants that are often not covered by SNP chips in appropriate proportions (Eynard et al., 2015). Previous work has shown that imposing a strict MAF filter may significantly affect the estimate of relatedness coefficients (Eynard et al. 2015), measures of genomic differentiation among populations (*F_ST_*, Hendricks et al., 2018) and may lead to inaccurate demographic inference (Nielsen et al., 2011). Removing rare alleles from data sets may also impede our ability to detect fine scale patterns of connectivity and local adaptation (O’Leary, Puritz, Willis, Hollenbeck, & Portnoy, 2018). Our results suggest that the MAF threshold has a direct impact on the GRM structure: applying a stringent MAF filter led to higher, and probably more accurate, pairwise relatedness coefficients among the most closely related individuals. More broadly, more stringent MAF filtering increased the variance in relatedness of the GRM, probably because most remaining genotyping errors were eliminated. However, the MAF threshold also had an impact on the *h^2^* estimate: *h^2^* tended to decrease when increasing the MAF threshold, even when the number of SNPs used to build the GRMs was fixed. This suggests that rare segregating variants of functional importance were excluded by mistake in an attempt to remove genotyping errors. This is in agreement with the recent work of Maroulie et al. (2017), who showed that low-frequency variants may individually have greater influence than common variants on adult height in humans.

Together, our results suggest that it is preferable to start with a high sequencing depth and the corresponding SNP calling procedure to obtain a high quality genomic dataset with few missing data rather than applying a stringent filtering procedure to offset low starting coverage (Catchen et al., 2013). A primary reason is that strict MAF or LM filters may lead to a sharp decline in the number of markers and so in our ability to capture the genetic variance of traits. The second is that it is particularly difficult to discriminate sequencing errors from low-frequency variants that may contribute to trait heritability.

### 4.3. Limits

While encouraging, our study also identified several limits to this approach. A first concern is that while the number of samples used here appears sufficient to detect high heritability (*h^2^*>0.50), it is probably not adequate to detect low heritability values or to estimate genetic covariance between traits (Perrier et al., 2018). We performed a power analysis using the GCTA-GREML power calculator (Visscher et al., 2014) (Fig S6). Given our number of samples (N=249) and the observed variance of the GRM (V_GRM_ = 1.5xl0^-3^), there is only a 50% chance of detecting a heritability that is lower or equal to 0.25. In our case, a minimum of 500 samples would be required to generate a power of nearly 100% for detecting values of heritability this low. Another important issue is that the variance of the GRM is inversely proportional to the sampling variance of the estimate of *h^2^* (Visscher & Goddard, 2015). Scientists must design their sampling scheme to capture the maximum variance in relatedness in their study population which is partly dependent on the social and spatial structure of the population (Flanagan et al., 2018). In our case study, the GRM mainly included distantly related individuals, reflecting the complex dynamics of the study population which has a high turn-over due to heavy hunting pressure and high dispersal rates. The GRM also captured family structure generated by highly related individuals establishing their home range close to each other within sampling sites. Values for V_GRM_ (1.5 x 10^-3^) are within the same range as those reported in Soay sheep (1.3 x 10^-3^) (Bérénos et al., 2014) and blue tits (4 x 10^-3^) (Perrier C. pers. obs), but one hundred times higher than between unrelated humans (Vinkhuyzen, Wray, Yang, Goddard, & Visscher, 2013) (2 x 10^-5^).

Another caveat is the extent of linkage disequilibrium (LD) that dictates how SNP loci tag causal loci and, thus, capture the genetic variance (Bérénos et al., 2014). Since the study population colonized the region only a few decades ago, we expected a low historical effective population size and, thus, a relatively high LD, as observed in Soay sheep (Bérénos et al., 2014). In accordance, we showed that a relatively modest number of markers (around 8000) appears to be enough to capture the genetic variance of the population. Hence, closely-linked SNPs that provide redundant information should be removed. However, without access to a reference genome sequence to physically map loci, we had no *a priori* knowledge of the genetic distance between SNPs. Estimating LD between each pair of SNPs could be an alternative option to prune statistically linked SNPs, however, this is computationally intensive. Moreover, pruning SNPs based solely on LD values, without any knowledge on their physical genomic distance, might be particularly risky since patterns of LD may also reflect the past demographic history of the population and/or effects of selection. Here, in order to reduce the aforementioned redundancy, we retained only one SNP per locus, but we did not filter for additional LD. Given the relatively modest number of RAD loci compared to the genome size, we also hypothesize a moderate representation of pairs of loci with high LD. Furthermore, Yang et al., (2015) showed that there is only a limited bias in *h^2^* due to heterogeneity in LD across the genome. Nevertheless, further analysis may be required to explore how *h^2^* estimates may be biased by LD among SNPs in our study system.

Lastly, a downside of pedigree-free estimation of quantitative genetic parameters is that partitioning of other variance components, such as maternal effects, is not possible without additional information. Here, heritabilities were estimated and interpreted for comparative purposes only, hence, we deliberately did not account for potential confounding environmental sources of similarity among individuals (e.g. maternal effect, habitat type). Hence, it is very likely that our *h^2^* estimates are biased upwards. This issue may be partly resolved by the recent development of methods to infer relationships between pairs of individuals (e.g. parent-offspring, full sibs, half-sibs) from SNP data (Andrews et al., 2018; Huisman, 2017).

### 4.4. Conclusion and prospects

Our study illustrates the huge potential of genomic-based relatedness for estimating quantitative genetic parameters in free-ranging populations (see also Bérénos et al., 2014; Malenfant, Davis, Richardson, Lunn, & Coltman, 2018; Perrier et al., 2018). Here, we showed that robust heritability estimates can be obtained from RAD-sequencing data in populations or species for which no genomic resources are available. This opens up new and exciting avenues in evolutionary biology, for example, by providing the opportunity to explore how the evolutionary potential of morphological, behavioural or life-history traits varies across space or time in virtually any species (Gienapp et al., 2017). This also paves the way towards more comparative/community quantitative genetic studies, for example, to explore how the heritability of traits varies among populations across environmental gradients (Martinez-Padilla et al 2017) or to better understand constraints on the evolution of phenotypic variation in several species interacting within a given ecosystem (Whitham et al., 2006).

## Supporting information

## ACKNOWLEDGMENTS

We thank numerous co-workers and volunteers for their assistance in field data collection, in particular Mark Hewison, Nicolas Morellet, J-M. Angibault, B. Cargnelutti, Y. Chaval, J-L. Rames, J. Joachim, B. Lourtet, D. Picot and N. Cebe. We also thank Karim Gharbi, Helene Gunter and others at the Edinburgh Genomics Facility for many useful discussions on RADseq and Andres Legarra, Isabel S. Winney, Caroline E. Thomson for their previous advices on the development of GRMs and implementation in animal models. We are grateful to the Genotoul Bioinformatics Plateform Toulouse Midi-Pyrénées (Genotoul Bioinfo) for providing help, computing and storage ressources. Comments from Mark Hewison, Nicolas Morellet helped improve the manuscript. This work was financially supported by the French National Institute for Agricultural Research (INRA), the UK Natural Environment Research Council (NERC) Biomolecular Analysis Facility at the University of Edinburgh (NBAF901), Centre National de la Recherche Scientifique (CNRS) core funding to BP, the EU within the framework of the Marie-Curie FP7 COFUND People Programme through the award of an Agreenskills fellowship to EQ, the Federal University of Toulouse through the PhD scholarship of LG (IDEX grant “GENEMOV” to BP and EQ).

## Data accessibility

Data will be available from the Dryad Digital Repository: http://dx.doi.org/xxxxx after final acceptance of the manuscript.

## Conflict of interest

The authors declare no conflict of interest

## Author Contributions

LG, EQ, BP, JP designed the study and interpreted the data; JM and EQ carried out the DNA extraction and quality controls; LG and MB performed the analyses; LG and EQ led the writing, all authors contributed to the manuscript and gave final approval for publication.

