## Supplementary material for "RAD-sequencing for estimating GRM-based heritability in the wild: a case study in roe deer"

**Table of Contents:**

| **Figure S1**. Map of the study area | Page 2 |
| --- | --- |
| **Figure S2**. Effect of the different sets of parameters in *Stacks* on allele  and locus error rates for the Low-coverage dataset | Page 2 |
| **Figure S3**. Biplot of genome wide relatedness matrices (GRM) | Page 3 |
| **Figure S4**. Sensitivity of h² estimates for body mass to SNP calling  (S*tacks* model) and Loci Missingness (LM) thresholds | Page 3 |
| **Figure S5***.* Estimated heritability of body mass as a function of  increasing number of SNPs | Page 4 |
| **Figure S6**. Statistical power for detecting heritability in our case study using  varying sample sizes | Page 4 |
| **Text S1**. Exploratory analysis of Stacks key assembly parameters  and SNP calling model using replicates | Page 5 |

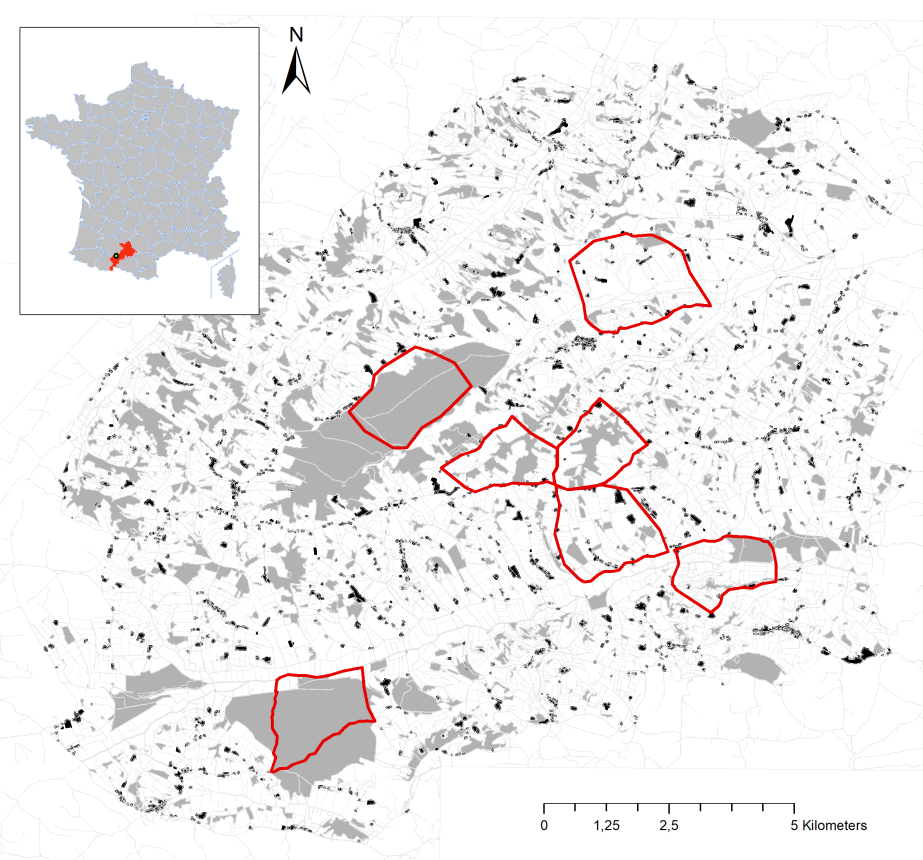

**Figure S1. Map of the study area.** Dwellings in black, woodland in grey, open areas in white. The boundaries of the seven sampling sites are delimited in red.

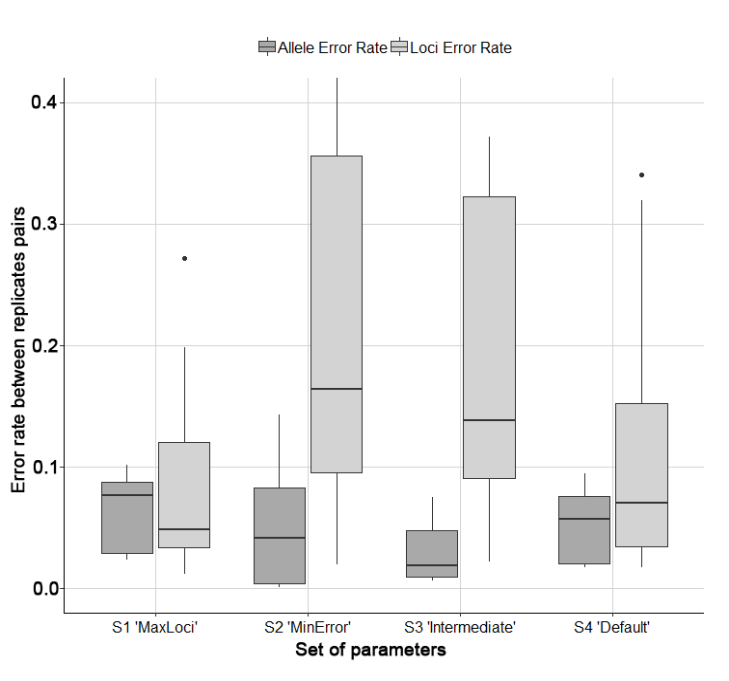

Figure S2. Effect of the different sets of parameters in *Stacks* on allele and locus error rates for the Low-coverage dataset (error rate distribution across the 7 replicate pairs). For each *Stacks* profile and each pair of technical replicates, we computed the locus error rate (LE_R_) corresponding to the number of loci present in only one of the two replicates, divided by the total number of loci being compared (i.e. missing data at the locus level) and the allele error rate (AE_R_), calculated as the number of incongruent genotypes between the two replicates, divided by the number of common loci.

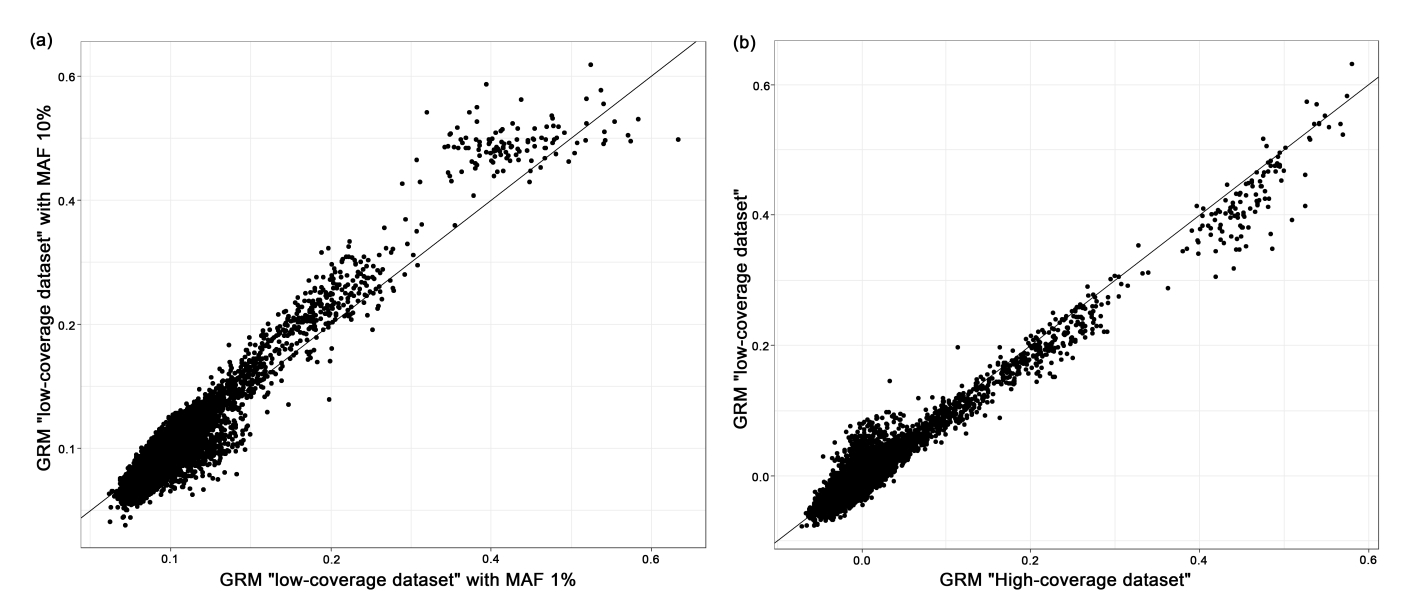

Figure S3. Biplot of genome wide relatedness matrices (GRM): (a) Genomic relatedness after removing loci with MAF<1% *versus* MAF<10% from the low-coverage SNP dataset, (b) Genomic relatedness using a MAF 1% filtering on the High *versus* Low coverage dataset.
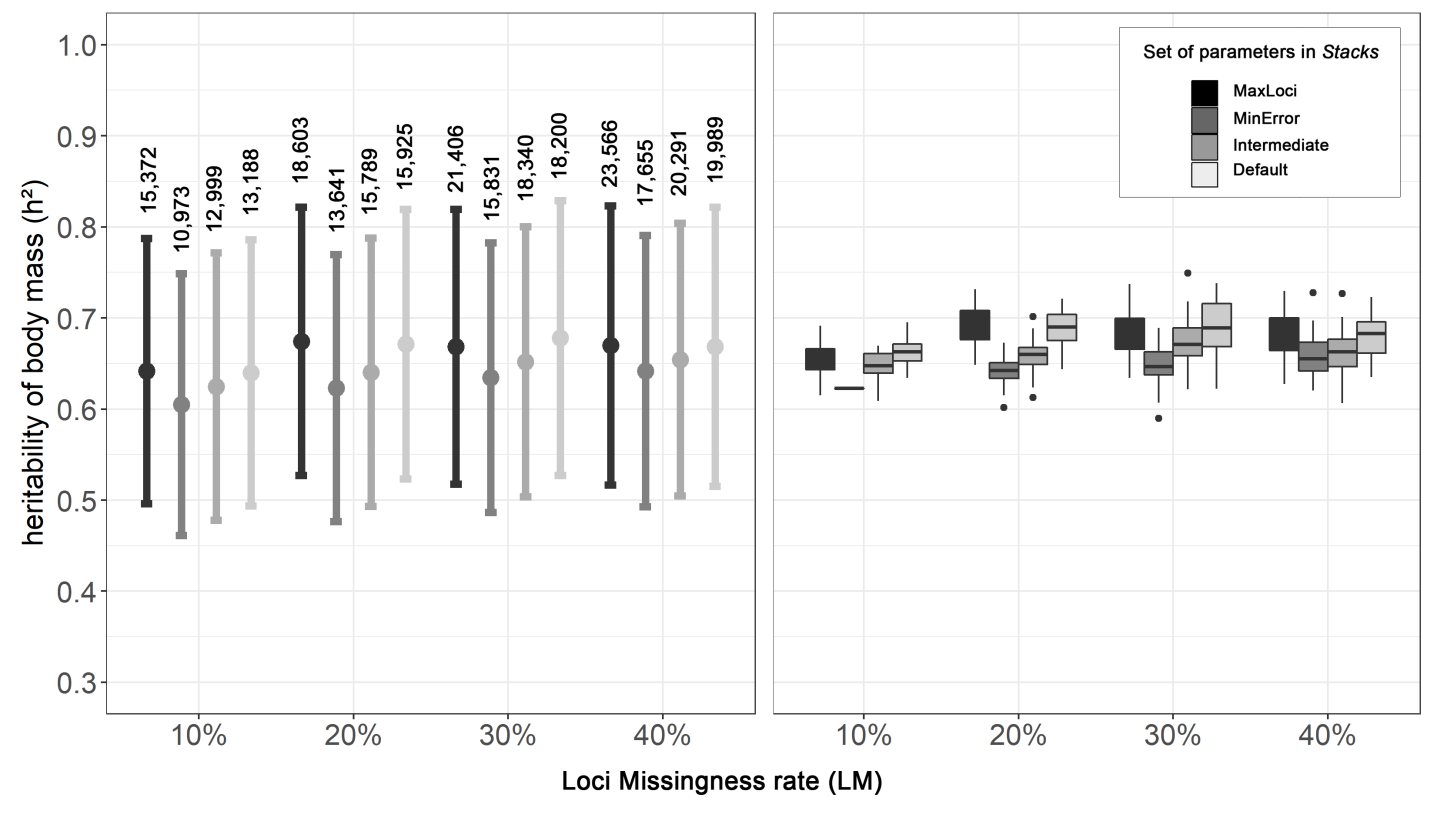

**Figure S4. Sensitivity of h² estimates for body mass to SNP calling (S*tacks* model) and Loci Missingness (LM) thresholds**: (a) raw *h²* estimates with standard error (error bars) obtained using GRMs computed with all SNPs recovered by a given SNP calling (Stacks model) and filtering method (LM = 10%, 20%, 30% or 30% and MAX=1%) (b) *h²* estimates obtained by fixing the number of SNPs to 10973 SNPs for all GRMs and resampling the SNP data 50 times. Boxplot and whiskers indicate average *h²* and variation across the 50 resampled datasets.

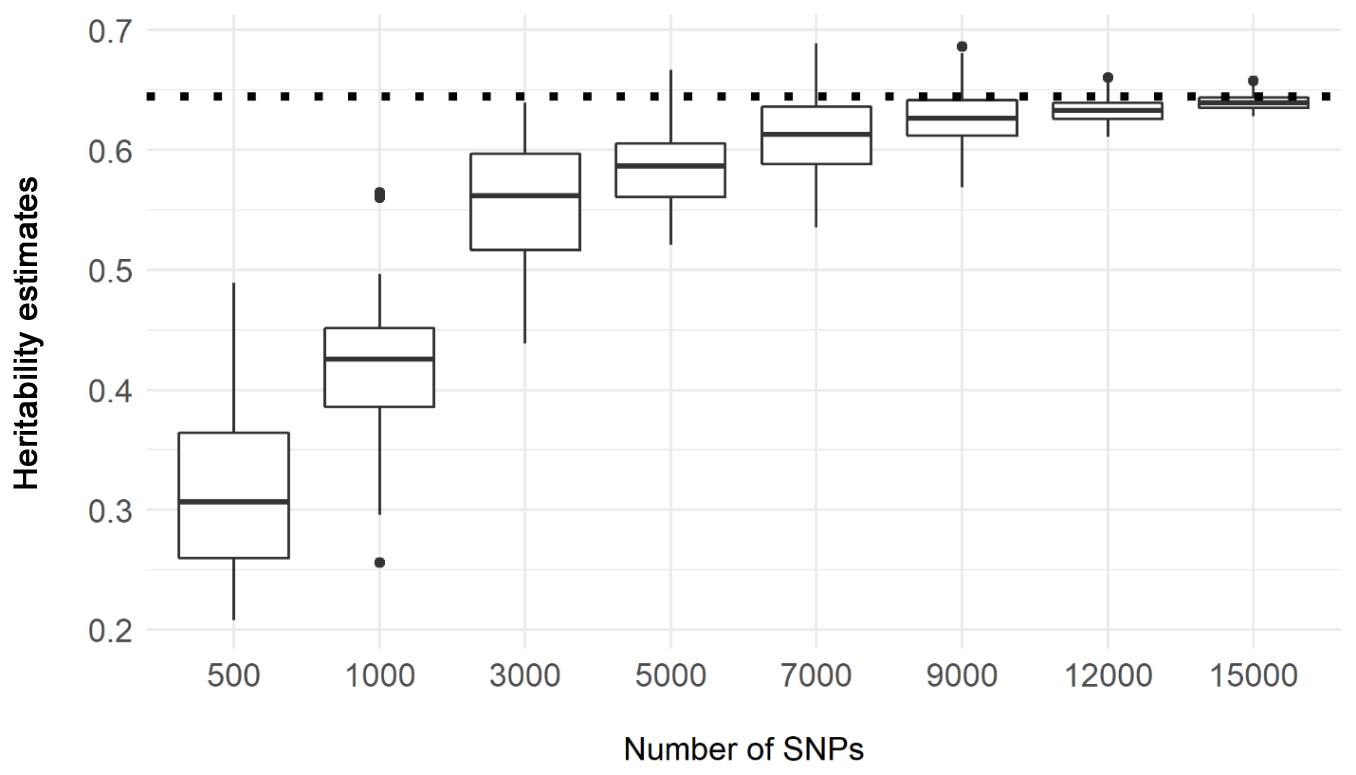

Figure S5*.* Estimated heritability of body mass as a function of increasing number of SNPs. Boxes and whiskers indicate the median and spread of 50 replicate sampled sets of SNPs. The dashed line represents the heritability estimates using all available markers. When less than 7000 markers were used to compute the GRM, heritability estimates are downward biased.

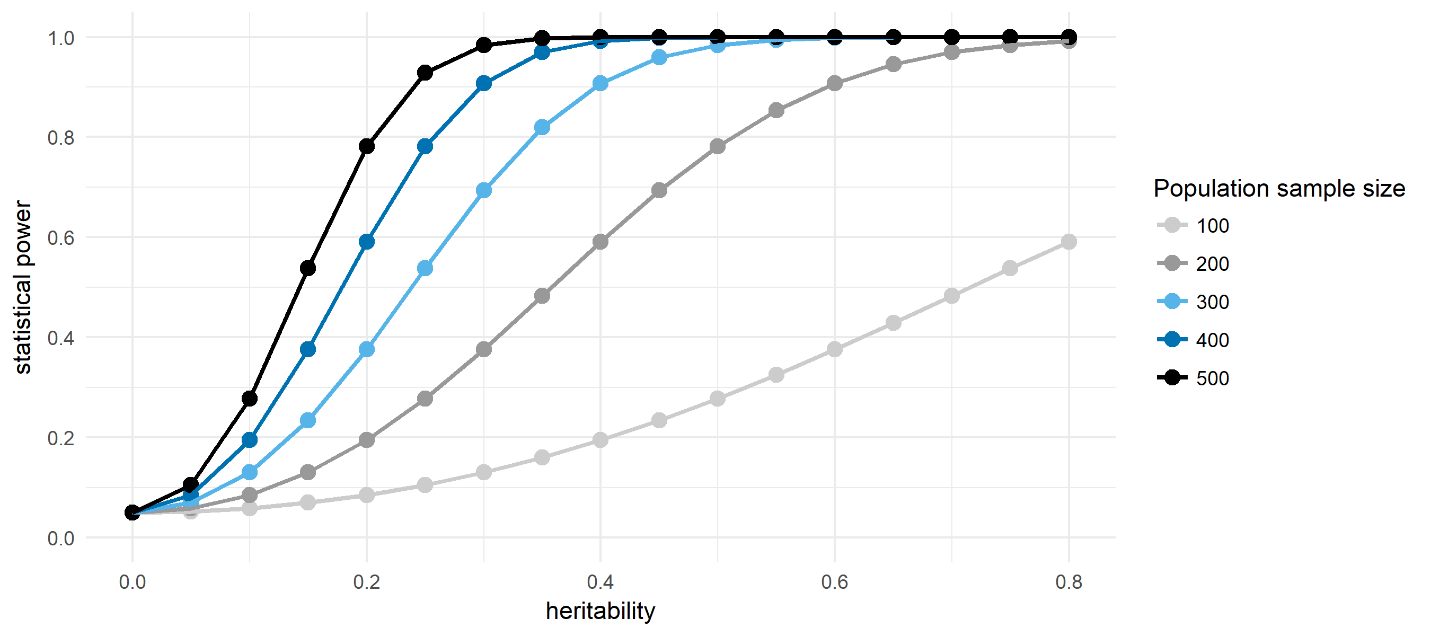

**Figure S6. Statistical power for detecting heritability in our case study using varying sample sizes.** Statistical power is the probability of detecting h²>0 giving a type I error rate of 0.05 for SNP-heritability going from 0 to 0.8 and populations sample size varying from 100 to 500 individuals. The power calculation was performed using the power calculator provided by Visscher et al. 2014 (Plos Genetics) using a variance in relatedness V_GRM_ of 1.5x10^-3^**.**

**Text S1. Exploratory analysis of Stacks key assembly parameters and SNP calling model using replicates.** Following the procedure and R scripts provided by Mastretta-Yanes 2015 in *Mol. Ecol*., we explored the effect of using different *de novo* assembly conditions and SNP calling model settings within *Stacks* on error rate and number of loci recovered. We used 7 pairs of replicates and ran *Stacks* multiple times with a range of parameter values. The four key parameters were tested with the range of values specified in parentheses: *(–m*) (*–M*) *(-n*) (*–max-locus_stacks)*. (-m) is the minimum number of raw reads required to form a stack (–m 2 to 15), the maximum number of mismatches allowed between stacks when processing an individual (-M 2 to 10), the allowed number of mismatches between loci when building the catalog (-n 0 to 5) and the maximum number of stacks per locus (--max_locus_stacks 2 to 6). Only one parameter was varied at a time while keeping the other parameters fixed to m=3, M=2, n=1 and max_locus_stacks=3. We retained one SNP per fragment and only analyzed SNPS that were present in >80% of samples. The outputs of this sensitivity analysis (see Fig A,B,C below) allowed us to define four *Stacks* models (sets of parameters) corresponding to different ways of dealing with the trade-off between data quantity and quality (see main text).

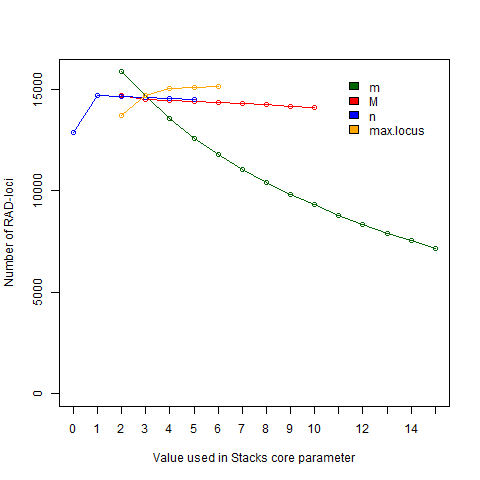

Fig_A- Total number of SNPs obtained using varying stacks code parameter settings. For each run, only one parameter was varied, with the others set to m=3, M= 2, n=0 and max_locus = 3.

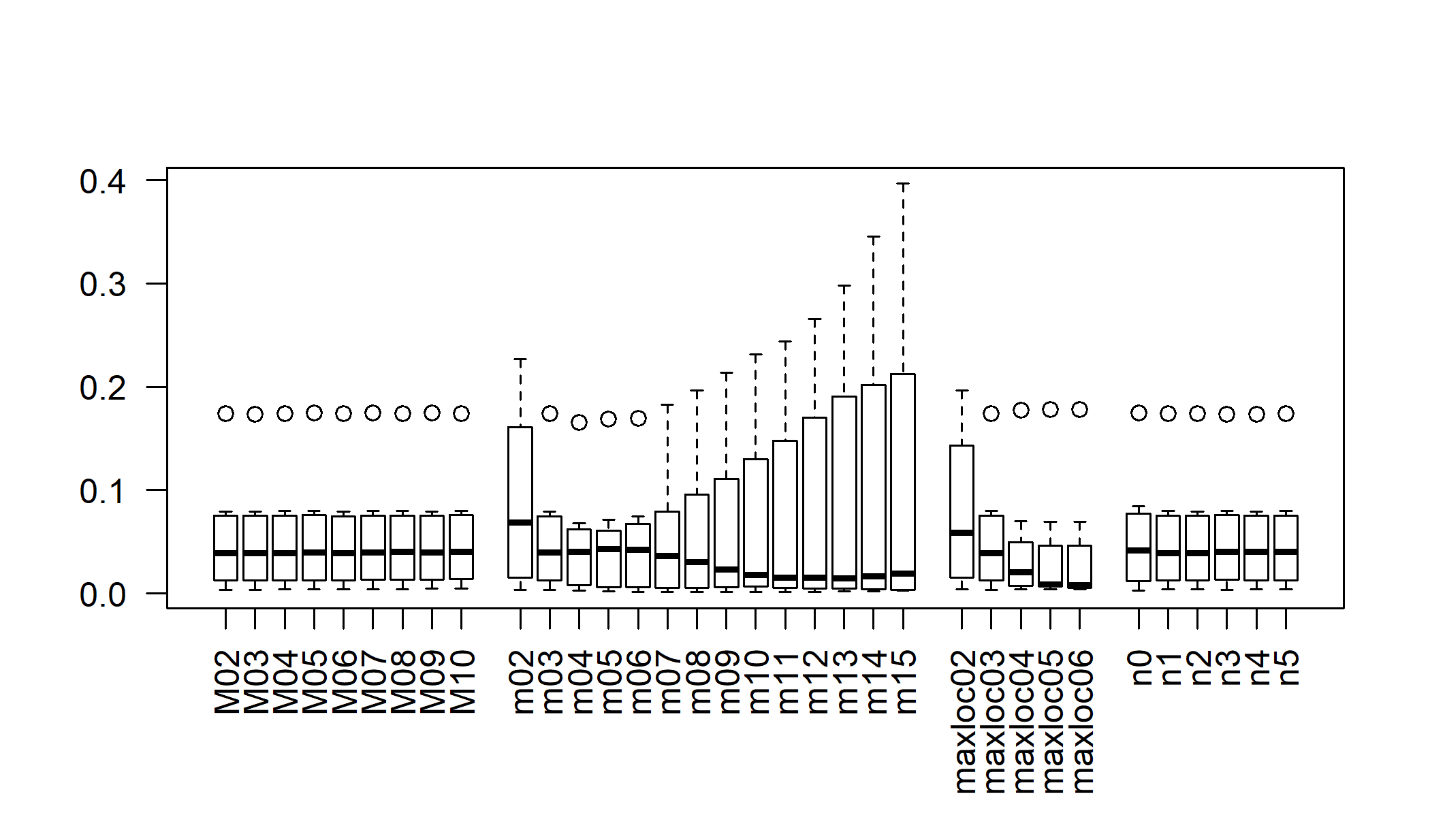

Fig_B- Effect of varying stacks parameter values on Loci error rate (LE_R_). For each run, only one parameter was varied (shown on the x axis), with settings for the others as explained in Fig. A.

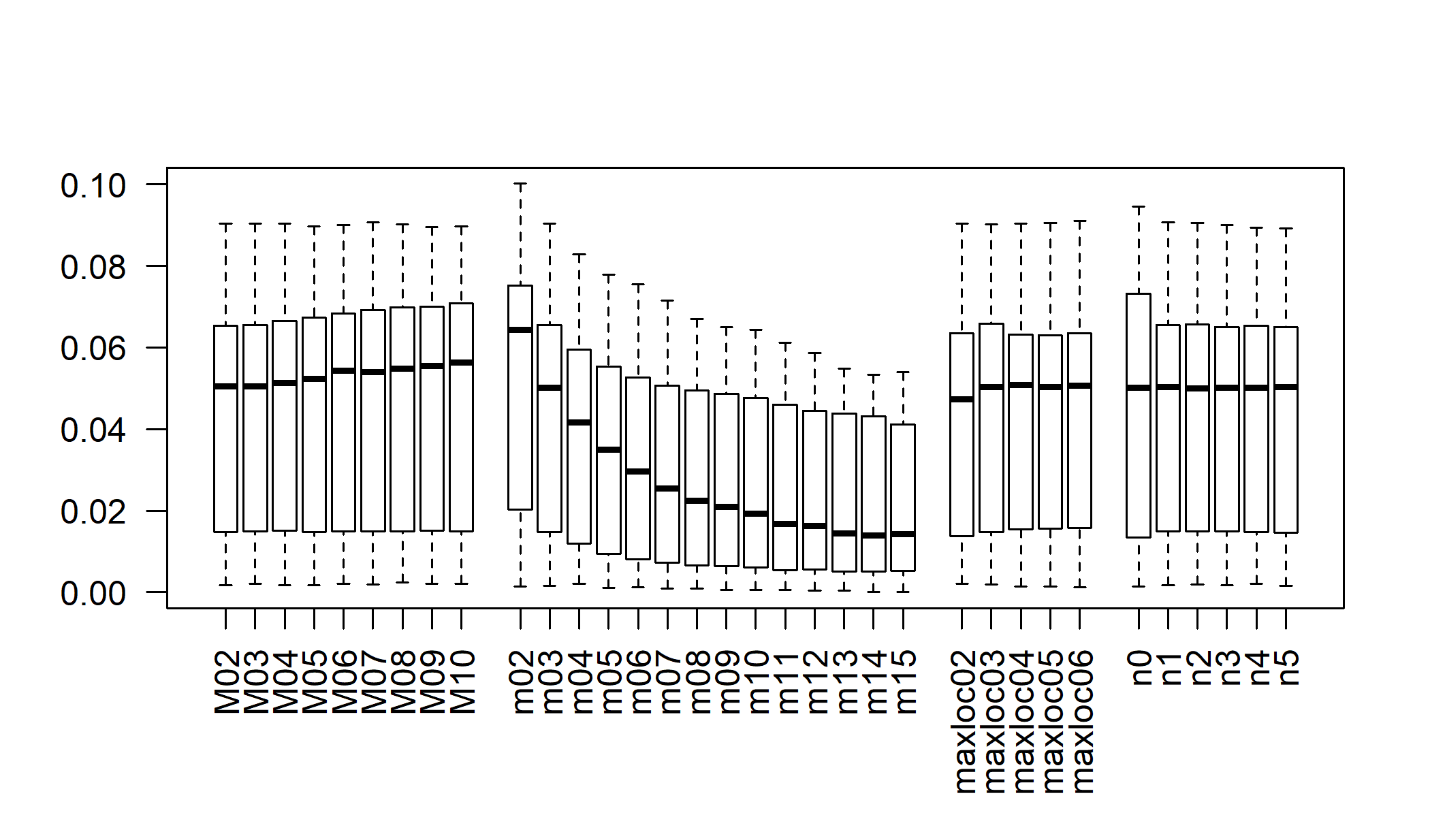

Fig_C- Effect of varying stacks parameter values on Allele error rate (AE_R_). For each run, only one parameter was varied (shown on the x axis), with settings for the others as explained in Fig. A.
